# Cell “hashing” with barcoded antibodies enables multiplexing and doublet detection for single cell genomics

**DOI:** 10.1101/237693

**Authors:** Marlon Stoeckius, Shiwei Zheng, Brian Houck-Loomis, Stephanie Hao, Bertrand Z. Yeung, Peter Smibert, Rahul Satija

## Abstract

Despite rapid developments in single cell sequencing technology, sample-specific batch effects, detection of cell doublets, and the cost of generating massive datasets remain outstanding challenges. Here, we introduce cell “hashing”, where oligo-tagged antibodies against ubiquitously expressed surface proteins are used to uniquely label cells from distinct samples, which can be subsequently pooled. By sequencing these tags alongside the cellular transcriptome, we can assign each cell to its sample of origin, and robustly identify doublets originating from multiple samples. We demonstrate our approach by pooling eight human PBMC samples on a single run of the 10x Chromium system, substantially reducing our per-cell costs for library generation. Cell “hashing” is inspired by, and complementary to, elegant multiplexing strategies based on genetic variation, which we also leverage to validate our results. We therefore envision that our approach will help to generalize the benefits of single cell multiplexing to diverse samples and experimental designs.

## INTRODUCTION

Single cell genomics offers enormous promise to transform our understanding of heterogeneous processes and to reconstruct unsupervised taxonomies of cell types^1,2^. As studies have progressed to profiling complex human tissues^3,4^ and even entire organisms^5,6^, there is a growing appreciation of the need for massively parallel technologies and datasets to uncover rare and subtle cell states^7-9^. While the per-cell cost of library prep has dropped, routine profiling of tens to hundreds of thousands of cells remains costly both for individual labs, and for consortia such as the Human Cell Atlas^10^.

Broadly related challenges also remain, including the robust identification of artifactual signals arising from cell doublets or technology-dependent batch effects^11^. In particular, reliably identifying expression profiles corresponding to more than one cell (hereby referred to as ‘multiplets’) remains an unsolved challenged in single cell RNA-seq (scRNA-seq) analysis, and a robust solution would simultaneously improve data quality and enable increased experimental throughput. While multiplets are expected to generate higher complexity libraries compared to singlets, the strength of this signal is not sufficient for unambiguous identification^11^. Similarly, technical and ‘batch’ effects have been demonstrated to mask biological signal in the integrated analysis of scRNA-seq experiments^12^, necessitating experimental solutions to mitigate these challenges.

Recent developments have poignantly demonstrated how sample multiplexing can simultaneously overcome multiple challenges^13,14^. For example, the demuxlet^13^ algorithm enables the pooling of samples with distinct genotypes together into a single scRNA-seq experiment. Here, the sample-specific genetic polymorphisms serve as a fingerprint for the sample of origin, and therefore can be used to assign each cell to an individual after sequencing. This workflow also enables the detection of multiplets originating from two individuals, reducing non-identifiable multiplets at a rate that is directly proportional to the number of multiplexed samples. While this elegant approach requires pooled samples to originate from previously genotyped individuals, in principle any approach assigning sample fingerprints that can be measured alongside scRNA-seq would enable a similar strategy. For instance, sample multiplexing is frequently utilized in flow and mass cytometry by labeling distinct samples with antibodies to the same ubiquitously expressed surface protein, but conjugated to different fluorophores or isotopes, respectively^15,16^.

We recently introduced CITE-seq^17^, where oligonucleotide-tagged antibodies are used to convert the detection of cell-surface proteins into a sequenceable read-out alongside scRNA-seq. We reasoned that a defined set of oligo-tagged antibodies against ubiquitous surface proteins could uniquely label different experimental samples. This enables us to pool these together, and use the barcoded antibody signal as a fingerprint for reliable demultiplexing. We refer to this approach as cell “hashing”, as our set of oligos defines a “look up table” to assign each multiplexed cell to its original sample. We demonstrate this approach by labeling and pooling eight human PBMC samples, and running them simultaneously in a single droplet based scRNA-seq run. Cell hashtags allow for robust sample multiplexing, confident multiplet identification, and the discrimination of low-quality cells from ambient RNA. In addition to enabling ‘super-loading’ of commercial scRNA-seq platforms to substantially reduce costs, this strategy represents a generalizable approach for doublet identification and multiplexing that can be tailored to any biological sample or experimental design.

## RESULTS

### Hashtag-enabled demultiplexing based on ubiquitous surface protein expression

We sought to extend antibody-based multiplexing strategies^15,16^ to scRNA-seq using a modification of our CITE-seq method. We chose a set of monoclonal antibodies directed against ubiquitously and highly expressed immune surface markers (CD45, CD98, CD44, and CD11a), combined these antibodies into eight identical pools (pool A through H), and subsequently conjugated each pool to a distinct hashtag oligonucleotide (henceforth referred to as HTO, Figure 1A; Methods). The HTOs contain a unique 12-bp barcode that can be sequenced alongside the cellular transcriptome, with only minor modifications to standard scRNA-seq protocols. We utilized an improved and simplified conjugation chemistry compared to our previous approach using iEDDA click chemistry to covalently attach oligonucleotides to antibodies^18^ (Methods).

**Figure 1.**
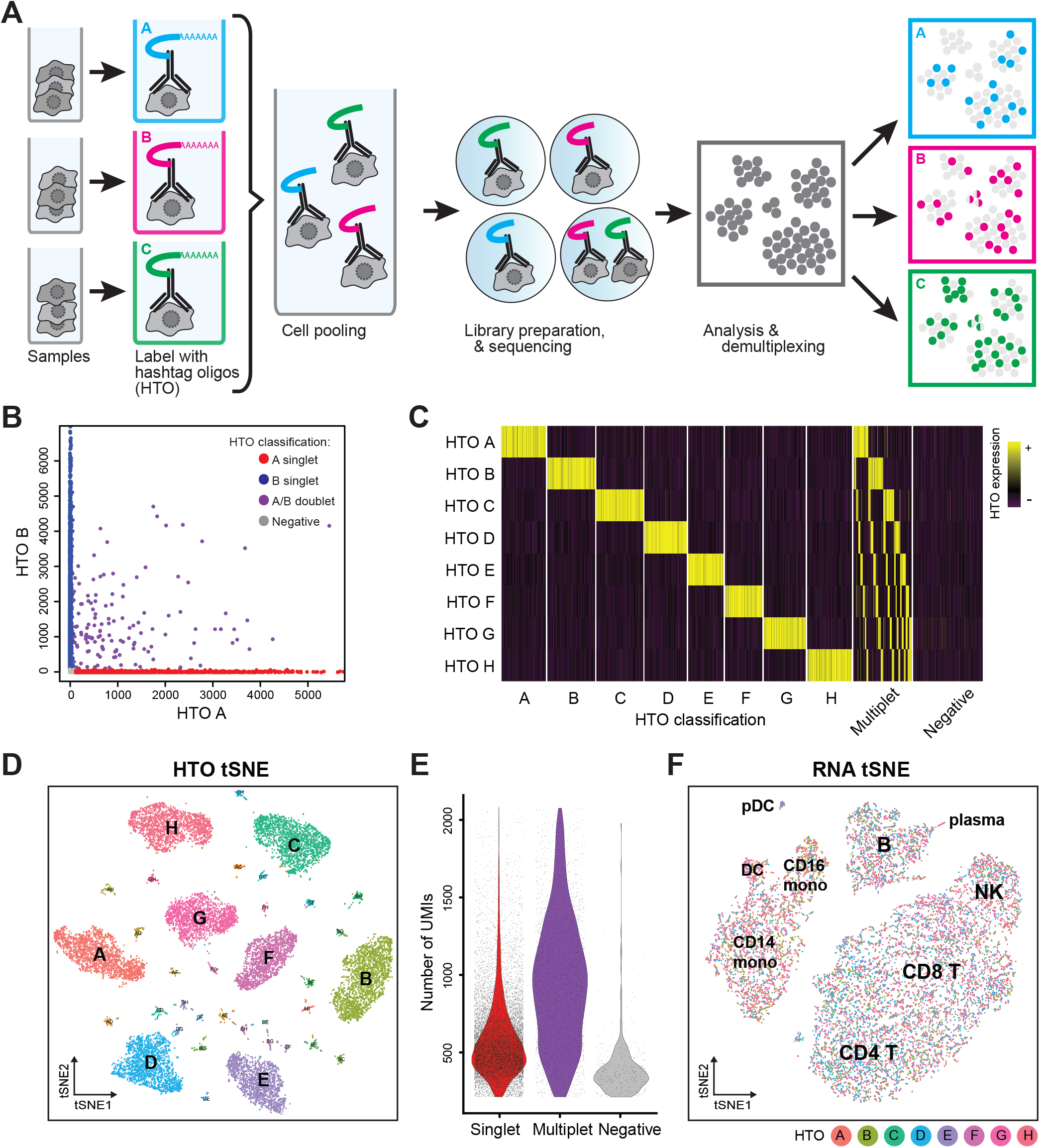
Sample multiplexing using DNA-barcoded antibodies. **A**) Schematic overview of sample multiplexing by cell hashing. Cells from different samples are incubated with DNA-barcoded antibodies recognizing ubiquitous cell surface proteins. Distinct barcodes (referred to as ‘hashtag’-oligos, HTO), on the antibodies allow pooling of multiple samples into one single cell RNA-sequencing experiment. After sequencing, cells can be classified to their sample of origin based on HTO levels (Methods). **B**) Representative scatter plot showing raw counts for HTO A and HTO B, across all cell barcodes. Both axes are clipped at 99.9% quantiles to exclude visual outliers. **C**) Heatmap of all normalized and scaled HTO levels, based on our classifications. Doublets and mulitplets express more than one HTO. Negative populations contain HEK-293T and mouse NIH-3T3 cells that were spiked into the experiments as negative controls. **D**) tSNE embedding of the HTO dataset. Cells are colored and labeled based on our classifications. Eight singlet clusters and all 28 cross-sample doublet clusters are clearly present. **E**) Distribution of RNA UMIs per cell barcode in cells that were characterized as singlets (red), doublets (violet) or negative (grey). **F**) Transcriptome-based clustering of single-cell expression profiles reveals distinct immune cell populations interspersed across donors. B, B cells; T, T cells; NK, natural killer cells; mono, monocytes; DC, dendritic cells; pDC, plasmacytoid dendritic cells; and plasma cells. Cells are colored based on their HTO classification (donor ID), as in (**D**).

We designed our strategy to enable CITE-seq and cell “hashing” to be performed simultaneously, but to generate separate sequencing libraries. Specifically, the HTOs contain a different amplification handle than our standard CITE-seq antibody derived tags (ADT) (Methods). This allows HTOs, ADTs, and scRNA-seq libraries to be independently amplified and pooled at desired quantities. Notably, we have previously observed robust recovery of antibody signals from highly expressed epitopes due to their extremely high copy number. This is in contrast to the extensive ‘drop-out’ levels observed for scRNA-seq data, and suggests that we can faithfully recover HTOs from each single cell, enabling assignment to sample of origin with high fidelity.

To benchmark our strategy and demonstrate its utility, we obtained PBMCs from eight separate human donors (referred to as donors A through H), and independently stained each sample with one of our HTO-conjugated antibody pools, while simultaneously performing a titration experiment with a pool of seven immunophenotypic markers (Methods) for CITE-seq. We subsequently pooled all cells together in equal proportion, alongside an equal number of unstained HEK-293T cells (and 3% mouse NIH-3T3 cells) as negative controls, and ran the pool in a single lane on the 10x Genomics Chromium Single Cell 3’ v2 system. Following the approach in Kang et al^13^, we ‘super-loaded’ the 10x Genomics instrument, loading cells at a significantly higher concentration with an expected yield of 20,000 single cells and 5,000 multiplets. Based on Poisson statistics, 4,365 multiplets should represent cell combinations from distinct samples and can potentially be discarded, leading to an unresolved multiplet rate of 3.1% Notably, achieving a similar multiplet rate without multiplexing would yield ~4,000 singlets. As the cost of commercial droplet-based systems is fixed per run, multiplexing therefore allows for the profiling of ~400% more cells for the same cost.

We performed partitioning and reverse transcription according standard protocols, utilizing only a slightly modified downstream amplification strategy (Methods) to generate transcriptome, HTO, and ADT libraries. We pooled and sequenced these on an Illumina HiSeq2500 (two rapid run flowcells), aiming for a 90%:5%:5% contribution of the three libraries in the sequencing data. Additionally, we performed genotyping of all eight PBMC samples and HEK-293T cells with the Illumina Infinium CoreExome array, allowing us to utilize both HTOs and sample genotypes (assessed by demuxlet^13^) as independent demultiplexing approaches.

When examining pairwise expression of two HTO counts, we observed relationships akin to ‘species-mixing’ plots (Figure 1B), suggesting mutual exclusivity of HTO signal between singlets. Extending beyond pairwise analysis, we developed a straightforward statistical model to classify each barcode as ‘positive’ or ‘negative’ for each HTO (Methods). Briefly, we modeled the ‘background’ signal for each HTO independently as a negative binomial distribution, estimating background cells based on the results of an initial k-medoids clustering of all HTO reads (Methods). Barcodes with HTO signals above the 99% quantile for this distribution were labeled as ‘positive’, and barcodes that were ‘positive’ for more than one HTO were labeled as multiplets. We classified all barcodes where we detected at least 200 RNA UMI, regardless of HTO signal.

Our classifications (visualized as a heatmap in Figure 1C) suggested clear identification of eight singlet populations, as well as multiplet groups. We also identified barcodes with negligible background signal for any of the HTOs (labeled as ‘negatives’), consisting primarily (87.5%) of HEK and mouse cells. We removed all HEK and mouse cells from downstream analyses (Methods), with the remaining barcodes representing 13,964 singlets, and 2,463 identifiable multiplets, in line with expectations. Our classifications were also fully concordant with a tSNE embedding, calculated using only the eight HTO signals, which enabled the clear visualization not only of the 8 groups of singlets (donors A through H), but also the 28 small groups representing all possible doublet combinations (Figure 1D). Moreover, we observed a clear positive shift in the distribution of RNA UMI/barcode for multiplets, as expected (Figure 1E), while the remaining negative barcodes expressed fewer UMIs and may represent failed reactions or ‘empty’ droplets containing only ambient RNA. These results strongly suggest that HTOs successfully assigned each barcode into its original sample, and enabled robust detection of cross-sample multiplets. Performing transcriptomic clustering of the classified singlets enabled clear detection of nine hematopoietic subpopulations, which were interspersed across all eight donors (Figure 1F).

### Genotype-based demultiplexing validates cell “hashing”

We next compared our HTO-based classifications to those obtained by demuxlet^13^. Overall we observed strong concordance between the techniques, even when considering the precise sample mixture in called doublets (Figure 2A). Exploring areas of disagreement, we identified 1,138 barcodes that were classified based on HTO levels as singlets, but were identified as ‘ambiguous’ by demuxlet. Notably, the strength of HTO classification for these discordant barcodes (represented by the number of reads assigned to the most highly expressed HTO) was identical to barcodes that were classified as singlets by both approaches (Figure 2B). However, discordant barcodes did have reduced RNA UMI counts (Figure 2C). We conclude that these barcodes likely could not be genetically classified at our shallow sequencing depth, which is below the recommended depth for using demuxlet, but likely represent true single cells based on our HTO classifications.

**Figure 2.**
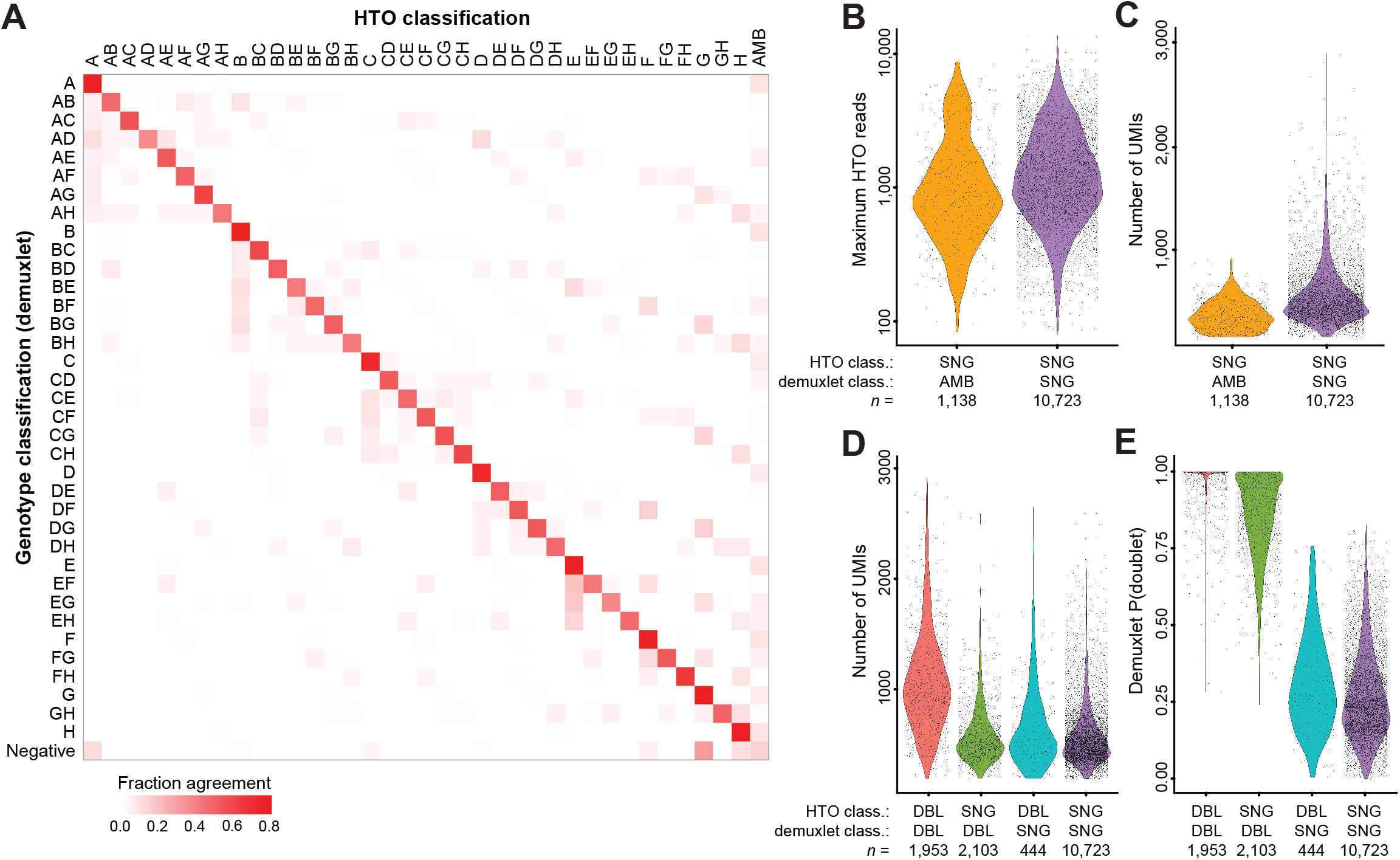
Validation of cell “hashing” using demuxlet. **A**) Row-normalized ‘confusion matrix’ comparing demuxlet and HTO classifications. Each value on the diagonal represents the fraction of barcodes for a given HTO classification that received an identical classification from demuxlet. **B**) Count distribution of the most highly expressed HTO for groups of concordant and discordant singlets. Both groups have identical classification strength based on cell “hashing”. **C**) Discordant singlets have lower UMI counts, suggesting that a lack of sequencing depth contributed to ‘ambiguous’ calls from demuxlet. **D**) RNA UMI distributions for discordant and concordant multiplets. Only concordant multiplets exhibit increased molecular complexity, suggesting that both methods are conservatively overcalling multiplets in discordant cases. **E**) In support of this, demuxlet assigns lower multiplets posterior probabilities to discordant calls.

In addition, we also observed 2,547 barcodes that received discordant singlet/doublet classifications between the two techniques (Figure 2D). We note that this does reflect a minority of barcodes (compared to 12,676 concordant classifications), and that in these discordant cases it is difficult to be certain which of these methods is correct. However, when we examined the UMI distributions of each classification group, we observed that only barcodes classified as doublets by both techniques exhibited a positive shift in transcriptomic complexity (Figure 2D). This suggests that these discordant calls are largely made up of true singlets, but represent conservative false positives from both methods, perhaps due to ambient RNA or HTO signal. Consistent with this interpretation, when we restricted our analysis to cases where demuxlet called barcodes as doublets with >95% probability, we observed a 71% drop in the number of discordant calls (Figure 2E).

### Cell hashing enables the efficient optimization of CITE-seq antibody panels

Our multiplexing strategy not only enables pooling across donors, but also the simultaneous profiling of multiple experimental conditions. This is widely applicable for simultaneous profiling of diverse environmental and genetic perturbations, but we reasoned that we could also efficiently optimize experimental workflows, such as the titration of antibody concentrations for CITE-seq experiments. In flow cytometry, antibodies are typically run individually over a large dilution series to assess signal to noise ratios and identify optimal concentrations^19^. While such experiments would be extremely cost prohibitive if run as individual 10x Genomics lanes, we reasoned that we could multiplex these experiments together using cell “hashing”.

We therefore incubated the PBMCs from different donors with a dilution series of antibody concentrations ranging over three orders of magnitude (Methods). Concentrations of CITE-seq antibodies were staggered between the different samples to keep the total amount of antibody and oligo consistent in each sample. After sample demultiplexing, we examined ADT distributions across all concentrations for each antibody (examples in Figure 3A-C), and assessed signal-to-noise ratio by calculating a staining index similar to commonly used metrics for flow cytometry optimization (Figure 3D) (Methods).

**Figure 3.**
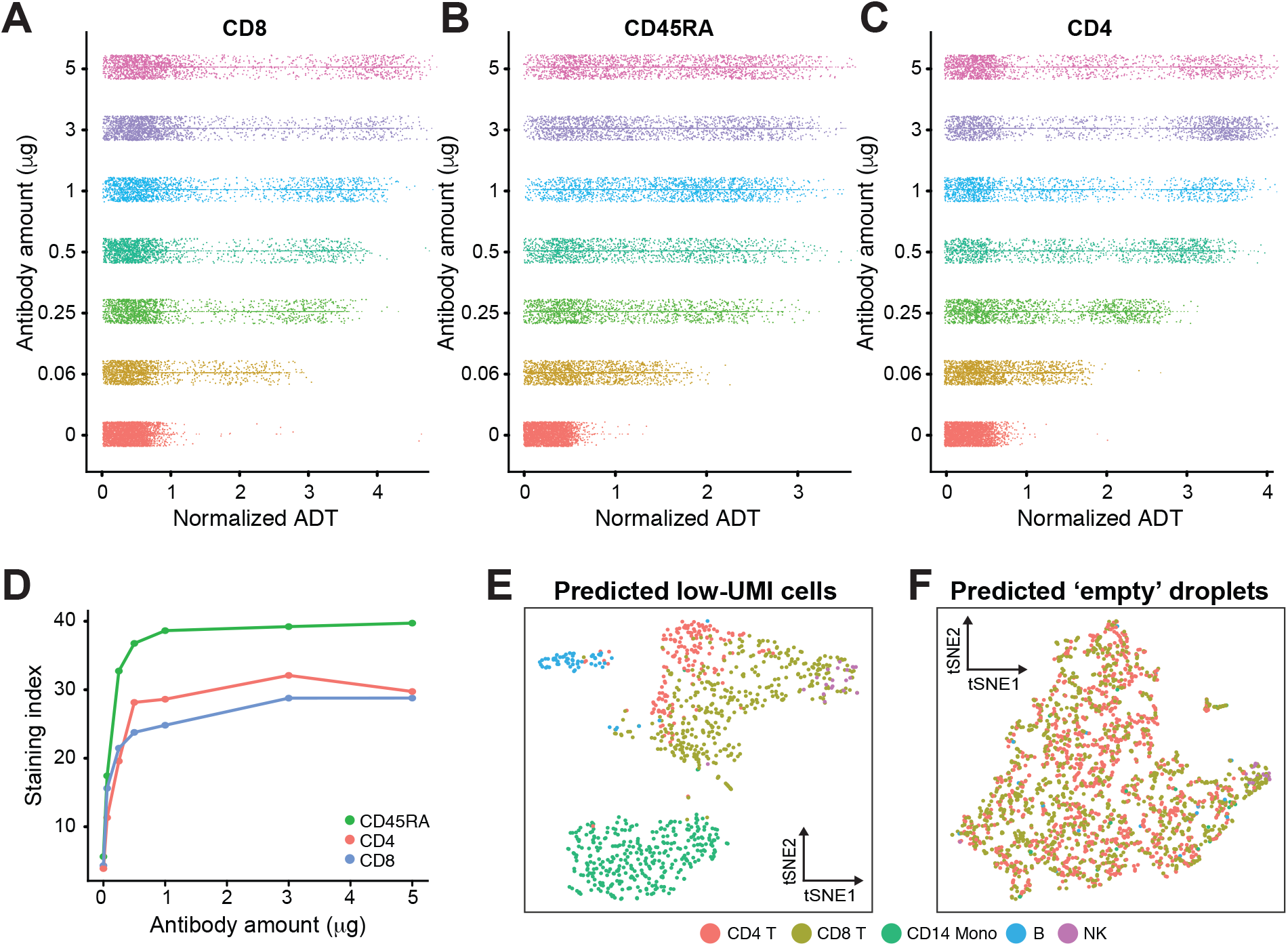
Cell “hashing” enables efficient experimental optimization and identification of low quality cells. **A-C**) We performed a titration series to assess optimal staining concentrations for a panel of CITE-seq immunophenotyping antibodies. Normalized ADT counts for CD8 (**A**), CD45RA (**B**) and CD4 (**C**) are depicted for the different concentrations used per test. D) Titration curve, depicting the staining index (SI; Methods) for these three antibodies across the titration series. The signal/noise ratio for these antibodies begins to saturate at levels similar to manufacturer recommended staining concentrations typical for flow cytometry antibodies. **E**) Cells with low UMI counts can be distinguished from ambient RNA using HTO classifications. Classified singlets group into canonical hematopoietic populations. **F**) Barcodes classified as ‘negative’ do not group into clusters, and likely represent ‘empty’ droplets containing only ambient RNA.

All antibodies exhibited only background signal in the negative control conditions, and very weak signal-to-noise at 0.06 μg/test. We observed that the signal-to-noise ratio for most antibodies began to saturate within the concentration range of 0.5 to 1 μg/test, comparable to the recommended concentrations for flow cytometry (Figure 3D). This experiment was meant as a proof-of-concept; an ideal titration experiment would use cells from the same donor for all conditions and a larger range of concentrations, but clearly demonstrates how cell “hashing” can be used to rapidly and efficiently optimize experimental workflows.

### Cell hashtags enable the discrimination of low quality cells from ambient RNA

Our cell hashtags can discriminate single cells from doublets based on the clear expression of a single HTO, and we next asked whether this feature could also distinguish low quality cells from ambient RNA. If so, this would enable us to reduce our UMI ‘cutoff’ (previously set at 200), and would allow for the possibility that certain barcodes representing ambient RNA may express more UMI than some true single cells. Most workflows set stringent UMI cutoffs to exclude all ambient RNA, biasing scRNA-seq results against cells with low RNA content, and likely skewing proportional estimates of cell type.

Indeed, when considering 3,473 barcodes containing 50-200 UMI, we recovered 954 additional singlets based on HTO classifications, with 2,432 barcodes characterized as negatives. We classified each barcode as one of our previously determined nine hematopoietic populations (Methods; Figure 1F), and visualized the results on a transcriptomic tSNE embedding, calculated independently for both ‘singlet’ and ‘negative’ groups. For predicted singlets, barcodes projected to B, NK, T, and myeloid populations which were consistently separated on tSNE, suggesting that these barcodes represent true single cells (Figure 3E). In contrast, ‘negative’ barcodes did not separate based on their forced classification, consistent with these barcodes reflecting ambient RNA mixtures that may blend multiple subpopulations. We therefore conclude that by providing a readout of sample identity that is independent of the transcriptome, cell “hashing” can help recover low quality cells that can otherwise be difficult to distinguish from ambient RNA (Figure 3F).

## DISCUSSION

Here, we introduce a new method for scRNA-seq multiplexing, where cells are labeled with sample-specific “hashtags” for downstream demultiplexing and doublet detection. Our approach is complementary to pioneering genetic multiplexing strategies, with each having unique advantages. Genetic multiplexing does not utilize exogenous barcodes, and therefore does not require alterations to existing workflows prior to or after sample pooling. In contrast, cell “hashing” requires incubation with antibodies against ubiquitously expressed surface proteins, but can multiplex samples with the same genotype. Both methods do slightly increase downstream sequencing costs, due to increased depth or read length needed to identify SNPs (genetic approaches), or sequencing of HTO libraries (cell “hashing”; approximately 5% of transcriptome sequencing costs). We believe that researchers will benefit from both approaches, enabling multiplexing for a broad range of experimental designs. In particular, we envision that our method will be most useful when processing genetically identical samples subjected to diverse perturbations (or experimental conditions/optimizations, as in our titration experiment), or to reduce the doublet rate when running cells from a single sample.

By enabling the robust identification of cell multiplets, both cell “hashing” and genetic multiplexing allow the “super loading” of scRNA-seq platforms. We demonstrate this in the context of the 10x Genomics Chromium system, but this benefit applies to any single cell technology that relies on Poisson loading for cell isolation. The per-cell cost savings for library preparation can therefore be significant, approaching an order of magnitude as the number of multiplexed samples increases. Notably, cell “hashing” enables even a single sample to be highly multiplexed, as cells can be split into an arbitrary number of pools. As clearly discussed in Kang et al^13^, savings in library prep are partially offset by reads originating from multiplets, which must be sequenced and discarded. Still, as sequencing costs continue to drop, and experimental designs seek to minimize technology-driven batch effects, multiplexing should facilitate the generation of large scRNA-seq and CITE-seq datasets. Informatic detection of multiplets based on transcriptomic data also remains an important challenge for the field, for example, to identify doublets originating from two cells within the same sample.

In our current study, we used a pool of antibodies directed against highly and ubiquitously expressed lymphocyte surface proteins as the vehicle for our HTOs. This strategy aimed to mitigate the possibility that stochastic or cell-type variation in expression of any one marker would introduce bias in HTO recovery. Going forward, we expect a more universal pool of antibodies directed against ubiquitously expressed markers to be used as a universal cell “hashing” reagent for studies beyond the hematopoietic system. With the increasing interest in single nucleus sequencing^20^, an additional set of “hashing” reagents directed against nuclear proteins would further generalize this approach. Beyond antibody/epitope interactions, cell or nuclei “hashing” could also be performed using any other means of attaching an oligo to a cell or nucleus, including other protein:protein interactions, aptamers^21^, or direct chemical conjugation of oligos to cells or nuclei. These improvements will further enable multiplexing strategies to generalize to diverse experiments regardless of species, tissue, or technology.

## DATA AVAILABILITY

Sequencing data is available from the Gene Expression Omnibus under accession GSE108313. Computational tools for classifying cells based on HTO levels will be released as part of the Seurat R package for single cell analysis, available at http://www.satijalab.org/seurat

## ACKNOWLEDGEMENTS

We acknowledge members of the NYGC Technology Innovation and Satija labs for critical discussions and support. We thank M. Coppo, S. Fennessey, B. Baysa and S. Pescatore at NYGC for sequencing support. This work was supported by the Chan Zuckerberg Initiative (HCA-A-1704-01895, to RS and PS), NIHR21-HG-009748 (to PS), and an NIH New Innovator Award (DP2-HG-009623, to RS).

## METHODS

### PBMC genotyping

Peripheral blood mononuclear cells were obtained from AllCells (USA). Genomic DNA was purified using the All-prep kit (Qiagen, USA) and genotyped using the Infinium core exome 24 array (Illumina, USA) according to manufacturer’s instructions.

### Cell culture

HEK293T (human) and NIH-3T3 (mouse) cells were maintained according to standard procedures in Dulbecco’s Modified Eagle’s Medium (Thermo Fisher, USA) supplemented with 10% fetal bovine serum (Thermo Fisher, USA) at 37°C with 5% CO2.

### Antibody-oligo conjugates

Antibody-oligo conjugates directed against CD8 [clone: RPA-T8], CD45RA [clone: HI100], CD4 [clone: RPA-T4], HLA-DR [clone: L243], CD3 [clone: UCHT1], CCR7 [clone: G043H7] and PD-1 [clone: EH12.2H7] were provided by BioLegend (USA) containing 1-2 conjugated oligos per antibody on average.

Antibodies used for cell hashing were obtained as purified, unconjugated reagents from BioLegend (CD45 [clone: HI30], CD98 [clone: MEM-108], CD44 [clone: BJ18], and CD11a [clone: HI111]) and were covalently and irreversibly conjugated to HTOs by iEDDA-click chemistry as previously described^18^. In short, antibodies were washed into 1X borate buffered saline (50 mM borate, 150 mM NaCl pH 8.5) and concentrated to 1 mg/ml using an Amicon Ultra 0.5 ml 30 kDa MWCO centrifugal filter (Millipore). Methyltetrazine-PEG4-NHS ester (Click Chemistry Tools, USA) was dissolved in dry DMSO and added as a 30-fold excess to the antibody and allowed to react for 30 minutes at room temperature. Residual NHS groups were quenched by the addition of glycine and unreacted label was removed via centrifugal filtration. 5’-amine HTOs were ordered from Integrated DNA Technologies (USA) and reacted with a 20fold excess of *trans*-cyclooctene-PEG4-NHS (Click Chemistry Tools, USA) in 1X borate buffered saline supplemented with 20% DMSO for 30 minutes. Residual NHS groups were quenched by the addition of glycine and residual label was removed by desalting (Bio-Rad Micro Bio-Spin P6). Antibody-oligo conjugates were formed by mixing the appropriate labeled antibody and HTO and incubating at room temperature for at least 1 hour. Residual methyltetrazine groups on the antibody were quenched by the addition of *trans*-cyclooctene-PEG4-acid and unreacted oligo was removed centrifugal filtration using an Amicon Ultra 0.5 ml 50 kDa MWCO filter (Millipore, USA).

### Antibody Titration Series

To test optimal concentration of Antibody-Oligo conjugates provided by BioLegend (USA) per CITE-seq experiment, we tested 5 μg, 3 μg, 1 μg, 0.5 μg, 0.25 μg, 0.06 μg, and 0 μg for each conjugate. Titrations were staggered over the different batches to keep the total concentration of antibodies and oligos consistent between conditions (see table below)

|  | Donor<br>A | Donor<br>B | Donor<br>C | Donor<br>D | Donor<br>E | Donor<br>F | Donor<br>G | Donor<br>H |
| --- | --- | --- | --- | --- | --- | --- | --- | --- |
| CD8 | 5 | 3 | 1 | 0.5 | 0.25 | 0.06 | 0 | 0 |
| PD.1 | 3 | 1 | 0.5 | 0.25 | 0.06 | 0 | 5 | 0 |
| CCR7 | 1 | 0.5 | 0.25 | 0.06 | 0 | 5 | 3 | 0 |
| CD3 | 0.5 | 0.25 | 0.06 | 0 | 5 | 3 | 1 | 0 |
| HLA.DR | 0.25 | 0.06 | 0 | 5 | 3 | 1 | 0.5 | 0 |
| CD4 | 0.06 | 0 | 5 | 3 | 1 | 0.5 | 0.25 | 0 |
| CD45RA | 0 | 5 | 3 | 1 | 0.5 | 0.25 | 0.06 | 0 |

### Sample pooling

PBMCs from different donors were independently stained with one of our HTO-conjugated antibody pools and a pool of 7 immunophenotypic markers for CITE-seq at different amounts (see above). All eight PBMC samples were pooled at equal concentration, alongside unlabeled HEK293T and mouse 3T3 as negative controls (see table below) and loaded into the 10x Chromium instrument (see below).

| Sample in pool | Labeled<br>with<br>Hashtag # | HTO Barcode | # cells<br>loaded | Estimated<br>cell yield |
| --- | --- | --- | --- | --- |
| PBMC Donor A | A | AGGACCATCCAA | ~3.5 K | ~2.5 K |
| PBMC Donor B | B | ACATGTTACCGT | ~3.5 K | ~2.5 K |
| PBMC Donor C | C | AGCTTACTATCC | ~3.5 K | ~2.5 K |
| PBMC Donor D | D | TCGATAATGCGA | ~3.5 K | ~2.5 K |
| PBMC Donor E | E | GAGGCTGAGCTA | ~3.5 K | ~2.5 K |
| PBMC Donor F | F | GTGTGACGTATT | ~3.5 K | ~2.5 K |
| PBMC Donor G | G | ACTGTCTAACGG | ~3.5 K | ~2.5 K |
| PBMC Donor H | H | TATCACATCGGT | ~3.5 K | ~2.5 K |
| HEK293T | - | - | ~3.5 K | ~2.5 K |
| NIH-3T3 | - | - | ~0.8 K | ~0.5 K |

### CITE-seq on 10x Genomics instrument

Cells were ‘stained’ with hashtagging antibodies and CITE-seq antibodies as described for CITE-seq^17^. ‘Stained’ and washed cells were loaded into 10x Genomics single cell 3’ v2 workflow and processed according to manufacturer’s instructions up until the cDNA amplification step (10x Genomics, USA). 2 pmol of HTO and ADT additive oligonucleotides were spiked into the cDNA amplification PCR and cDNA was amplified according to the 10x Single Cell 3’ v2 protocol (10x Genomics, USA). Following PCR, 0.6X SPRI was used to separate the large cDNA fraction derived from cellular mRNAs (retained on beads) from the ADT- and hashtag-containing fraction (in supernatant). The cDNA fraction was processed according to the 10x Genomics Single Cell 3’ v2 protocol to generate the transcriptome library. An additional 1.4X reaction volume of SPRI beads was added to the ADT/hashtag fraction to bring the ratio up to 2.0X. Beads were washed with 80% ethanol, eluted in water, and an additional round of 2.0X SPRI performed to remove excess single stranded oligonucleotides from cDNA amplification. After final elution, separate PCRs were set up to generate the CITE-seq ADT library (SI-PCR and RPI-x primers), and the hashtag library (SI-PCR and D7xx_s). A detailed and regularly updated point-by-point protocol for CITE-seq, cell-hashtagging, and future updates can be found at www.cite-seq.com

### Oligonucleotide sequences

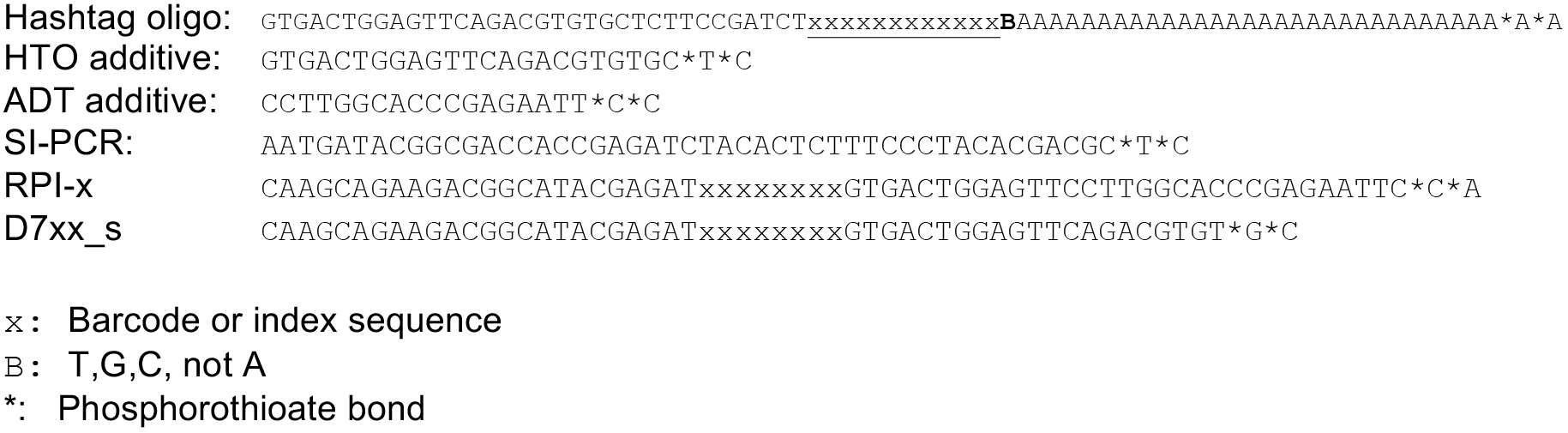

## COMPUTATIONAL METHODS

### Single-cell data processing

Fastq files from the 10x libraries with four distinct barcodes were pooled together and processed using the standard Drop-seq pipeline (Drop-seq tools v1.0, McCarroll Lab). Reads were aligned to the hg19-mm10 concatenated reference, and we included the top 50,000 cell barcodes in the raw digital expression matrix as output from Drop-seq tools. For ADT and HTO quantification, we implemented our previously developed tag quantification pipeline^17^ as a python script, available at https://github.com/Hoohm/CITE-seq-Count, and run with default parameters (maximum hamming distance of 1).

### Demultiplexing with genotyping data using demuxlet

We first generated a VCF file that contained the individual genotype (GT) from the Infinium core exome 24 array output, using the PLINK command line tools (version 1.07). This VCF file (which contained genotype information for the 8 PBMC donors as well as HEK cells), and the tagged bam file from Drop-seq pipeline were used as inputs for demuxlet^13^, with default parameters.

### Single-cell RNA data processing

Normalization and downstream analysis of RNA data were performed using the Seurat R package (version 2.1, Satija Lab) which enables the integrated processing of multi-modal (RNA, ADT, HTO) single cell datasets^22,23^. We collapsed the joint-species RNA expression matrix to only include the top 100 most highly expressed mouse genes (along with all human genes) using the CollapseSpeciesExpressionMatrix function.

We first considered a set of 22,119 barcodes where we detected at least 200 UMI in the transcriptome data. Since the HEK and 3T3 cells were not labeled with HTOs, we identified these cells based on their transcriptomes. We performed a low-resolution pre-clustering by performing PCA on the 500 most highly expressed genes, followed by Louvain-Jaccard clustering on a distance matrix based on the first five principal components^24-26^. Based on this clustering, we identified 248 3T3 cells and 3,401 HEK cells, with the remainder representing PBMCs.

As a separate test of HEK identity, we examined the demuxlet genotype for possible HEK cells. We observed 1,668 barcodes classified as HEK by the demuxlet algorithm, but whose transcriptomes clustered with PBMCs. These cells expressed ten-fold fewer UMI compared to transcriptomically-classified HEK cells, and did not express HEK-specific transcripts (i.e. NGFRAP1), both consistent with a PBMC identity. We therefore excluded these barcodes from all further analysis.

### Classification of barcodes based on HTO levels

HTO raw counts were normalized using centered log ratio (CLR) transformation, where counts were divided by the geometric mean of an HTO across cells, and log-transformed:

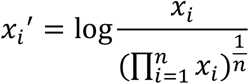

Here *x_i_* denotes the count for a specified HTO in cell *i*, *n* is the total cell number, *log* denotes the natural log. Pairwise analysis of either normalized or raw HTO counts (Figure 1B) revealed mutually exclusive relationships, though determining the exact cutoffs for positive and negative signals required further analysis. We reasoned that if we could determine a background distribution for each HTO based on ‘negative’ cells, outliers from this distribution would represent positive signals.

To assist in the unsupervised identification of ‘negative’ cells, we performed an initial k-medoids clustering for all cells based on the normalized HTO data. We set k=9, and observed (as expected) that eight of the clusters were highly enriched for expression of a particular HTO, while the ninth cluster was highly enriched for cells with low expression of all HTO. This represents an initial solution to the de-multiplexing problem that suggests likely populations of ‘positive’ and ‘negative’ cells for statistical analysis.

Following clustering, we performed the following the following procedure independently for each of the 8 HTOs. We identified the k-medoids cluster with the highest average HTO expression, and excluded these cells. We next fit a negative binomial distribution to the remaining HTO values, after further excluding the highest 0.5% values as potential outliers. We calculated the q=0.99 quantile of the fitted distribution, and thresholded each cell in the dataset based on this HTO-specific value.

We used this procedure to determine an ‘HTO classification’ for each barcode. Barcodes that were positive for only one HTO were classified as singlets. Barcodes that were positive for two or more HTO were classified as doublets, and assigned sample IDs based on their two most highly expressed HTO. Barcodes that were negative for all eight HTO were classified as ‘negative’.

We expect that barcodes classified as ‘singlets’ represent single cells, as we detect only a single HTO. However, they could also represent doublets of a PBMC with a HEK or 3T3 cell, as the latter two populations were unlabeled and represent negative controls. Indeed, when we analyzed the ‘HTO classification’ of cells that were transcriptomically annotated as HEK or 3T3 cells, we found that 73.4% were annotated as ‘negative’, while 29.2% were annotated as singlets, in complete agreement with expected ratios in our ‘super-loaded’ 10x experiment. These cells appear in the heatmap in Figure 1C, but all HEK and 3T3 cells were excluded from further analysis.

For two-dimensional visualization of HTO levels (Figure 1D), we used Euclidean distances calculated from the normalized HTO data as inputs for tSNE. Cells are colored based on their HTO classification as previously described. For visualization and clustering based on transcriptomic data (Figure 1F), we first performed PCA on the 2,000 most highly variable genes (as determined by variance/mean ratio), and used the distance matrix defined by the first 11 principal components as input to tSNE and graph-based clustering in Seurat (Figure 1E). We annotated the nine clusters based on canonical markers for known hematopoietic populations.

### Comparison with demuxlet

Demuxlet classifications were labeled as singlets (SNG), doublets (DBL) or ambiguous (AMB) according to the BEST column in the *.best output file. In Figure 2E, we plot the posterior probability of a doublet assignment, from the PRB.DBL column in the same file.

### Calculation of staining index for antibody titrations

To assess the optimal staining efficiency for CITE-seq experiments, we considered ADT levels for cells across a range of antibody concentrations, as multiplexed in a titration series. ADT levels were normalized using a CLR transformation of raw counts, using an identical approach to the normalization of HTO levels as previously described.

After normalization, we computed a staining index based on standard approaches in flow cytometry, which examine the difference between positive and negative peak medians, divided by the spread (i.e. twice the mean absolute deviation) of the negative peak.

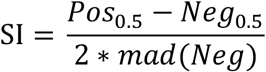

In order to avoid manual classification of positive and negative peaks, we implemented an automated procedure that can scale to multiple antibodies and concentrations. To approximate the negative peak, we leveraged unstained control cells (Donor H). To approximate the positive peak, we clustered the ADT data in each titration experiment (DonorA through DonorG). To perform clustering, we computed a Euclidean distance matrix across cells based on normalized ADT levels, and used this as input to the FindClusters function in Seurat with default parameters. We examined the results to identify the cluster with the maximally enriched ADT signal, and referred to the distribution of ADT levels within this cluster as the positive peak.

### Discriminating low-quality cells from ambient RNA

We performed HTO classification of low-quality barcodes (expressing between 50 and 200 UMI), using the previously determined HTO thresholds. For each barcode, we classified its expression as one of our previously determined nine hematopoietic populations using random forests, as implemented in the ranger package in R^27^. We first trained a classifier on the 13,757 PBMCs, using the 2,000 most variable genes as input, and their clustering identities as training labels. We then applied this classifier to each of the low-quality barcodes. We note that this classifier is guaranteed to return a result for each barcode.

## References

1. Stubbington, M. J. T., Rozenblatt-Rosen, O., Regev, A. & Teichmann, S. A. Single-cell transcriptomics to explore the immune system in health and disease. Science 358, 58–63 (2017).

2. Tanay, A. & Regev, A. Scaling single-cell genomics from phenomenology to mechanism. Nature 541, 331–338 (2017).

3. Villani, A.-C. et al. Single-cell RNA-seq reveals new types of human blood dendritic cells, monocytes, and progenitors. Science 356, (2017).

4. Velten, L. et al. Human haematopoietic stem cell lineage commitment is a continuous process. Nature Cell Biology 19, 271–281 (2017).

5. Cao, J. et al. Comprehensive single-cell transcriptional profiling of a multicellular organism. Science 357, 661–667 (2017).

6. Karaiskos, N. et al. The Drosophila embryo at single-cell transcriptome resolution. Science 8, eaan3235–14 (2017).

7. Macosko, E. Z. et al. Highly Parallel Genome-wide Expression Profiling of Individual Cells Using Nanoliter Droplets. Cell 161, 1202–1214 (2015).

8. Klein, A. M. et al. Droplet Barcoding for Single-Cell Transcriptomics Applied to Embryonic Stem Cells. Cell 161, 1187–1201 (2015).

9. Zheng, G. X. Y. et al. Massively parallel digital transcriptional profiling of single cells. Nature Communications 8, 1–12 (2017).

10. Regev, A. et al. Science Forum: The Human Cell Atlas. eLife 6, e27041 (2017).

11. Stegle, O., Teichmann, S. A. & Marioni, J. C. Computational and analytical challenges in single-cell transcriptomics. Nature Publishing Group 16, 133–145 (2015).

12. Hicks, S. C., Townes, F. W., Teng, M. & Irizarry, R. A. Missing data and technical variability in single-cell RNA-sequencing experiments. Biostatistics (2017). doi:10.1093/biostatistics/kxx053

13. Kang, H. M. et al. Multiplexed droplet single-cell RNA-sequencing using natural genetic variation. Nature Biotechnology (2017). doi:10.1038/nbt.4042

14. Tung, P.-Y. et al. Batch effects and the effective design of single-cell gene expression studies. Scientific Reports 7, 39921 (2017).

15. Krutzik, P. O. & Nolan, G. P. Fluorescent cell barcoding in flow cytometry allows high-throughput drug screening and signaling profiling. Nat Meth 3, 361–368 (2006).

16. Lai, L., Ong, R., Li, J. & Albani, S. A CD45-based barcoding approach to multiplex mass-cytometry (CyTOF). Cytometry 87, 369–374 (2015).

17. Stoeckius, M. et al. Simultaneous epitope and transcriptome measurement in single cells. Nat Meth 9, 2579–10 (2017).

18. van Buggenum, J. A. G. L. et al. A covalent and cleavable antibody-DNA conjugation strategy for sensitive protein detection via immuno-PCR. Nature Publishing Group 1–12 (2016). doi:10.1038/srep22675

19. Hulspas, R. Titration of fluorochrome-conjugated antibodies for labeling cell surface markers on live cells. Curr Protoc Cytom Chapter 6, Unit 6.29 (2010).

20. Lake, B. B. et al. A comparative strategy for single-nucleus and single-cell transcriptomes confirms accuracy in predicted cell-type expression from nuclear RNA. Scientific Reports 1–8 (2017). doi:10.1038/s41598017-04426-w

21. Delley, C. L., liu, L., Sarhan, M. F. & Abate, A. R. Combined aptamer and transcriptome sequencing of single cells. bioRxiv 1–10 (2017). doi:10.1101/228338

22. Satija, R., Farrell, J. A., Gennert, D., Schier, A. F. & Regev, A. Spatial reconstruction of single-cell gene expression data. Nature Biotechnology 33, 495–502 (2015).

23. Butler, A. & Satija, R. Integrated analysis of single cell transcriptomic data across conditions, technologies, and species. bioRxiv (2017). doi:10.1101/164889

24. Blondel, V. D., Guillaume, J.-L., Lambiotte, R. & Lefebvre, E. Fast unfolding of communities in large networks. J. Stat. Mech. 2008, P10008 (2008).

25. Levine, J. H. et al. Data-Driven Phenotypic Dissection of AML Reveals Progenitor-like Cells that Correlate with Prognosis. Cell 162, 184–197 (2015).

26. Shekhar, K. et al. Comprehensive Classification of Retinal Bipolar Neurons by Single-Cell Transcriptomics. Cell 166, 1308–1323. e30 (2016).

27. Wright, M. N. & Ziegler, A.ranger: A Fast Implementation of Random Forests for High Dimensional Data in C and R. Journal of Statistical Software 77, (2017).

